# Horizontal gene flow into Geobacillus is constrained by the chromosomal organization of growth and sporulation

**DOI:** 10.1101/381442

**Authors:** Alexander Esin, Tom Ellis, Tobias Warnecke

## Abstract

Horizontal gene transfer (HGT) in bacteria occurs in the context of adaptive genome architecture. As a consequence, some chromosomal neighbourhoods are likely more permissive to HGT than others. Here, we investigate the chromosomal topology of horizontal gene flow into a clade of Bacillaceae that includes Geobacillus *spp*. Reconstructing HGT patterns using a phylogenetic approach coupled to model-based reconciliation, we discover three large contiguous chromosomal zones of HGT enrichment. These zones encompass and connect classically defined genomic islands. Analyzing topological and strand biases of recent and older transfer events, we show that restrictions on entry are rapidly enforced by selection and that restrictive and permissive zones have existed in their current locations for long periods of evolution. The largest zone, characterized by a high influx of metabolic genes, is centred on the terminus. The other two zones flank a narrow non-permissive zone around the origin of replication and extend to delimit the first third of the chromosome – the part of the chromosome that is confined to the forespore during early spore formation. Horizontal transfers into this area are biased towards functions classically controlled by the forespore-specific sigma factor σ^F^: signal transduction, transcription, and particularly membrane biogenesis. Similar enrichment patterns are present in spore-forming but absent in non-spore-forming Bacilli. Our results suggest that the topology of HGT in Geobacillus, and Bacilli more generally, reflects constraints imposed by chromosomal organization for fast and sporulation, as asymmetric chromosomal entrapment in the forespore during early spore formation restricts where HGT-driven innovation in sporulation can occur.

## Introduction

Horizontal gene transfer (HGT) is ubiquitous amongst prokaryotes (Treangen and Rocha 2011) and frequently underpins the colonization of new niches via metabolic innovation (Hehemann et al. 2016; Gogarten and Townsend 2005; Pál et al. 2005). At the same time, most integration events are likely deleterious. Even if a gene arrives in one piece and manages to integrate into the host genome, a different genetic code, deviant codon usage patterns or promoter dependencies (Sorek et al. 2007; Moszer et al. 1999) can all undermine an easy *plug-and-play*, as can reliance on other molecular components. For example, genes whose protein products are highly connected, such as those that form part of large protein complexes, are less likely to be transferred than monomeric enzymes (Jain et al. 1999; Cohen et al. 2011). In short, new arrivals have to play nicely by themselves and with others. Often, they do neither.

This principle also extends to the chromosomal context of integration, with some neighbourhoods more and others less accommodating to newcomers. Bacterial genome organization can be highly stereotyped and location often matters for gene function and fitness. Highly expressed genes, for instance, are enriched around the origin of replication (*ori*), where increased gene dosage during replication transiently drives up expression (Couturier and Rocha 2006). Moving highly expressed native genes away from the origin is often detrimental. This has been demonstrated directly by repositioning experiments (Soler-Bistué et al. 2015) and is consistent with widespread conservation of origin-proximal locations for genes involved in translation and growth (Rocha 2008; Sobetzko et al. 2012; Repar and Warnecke 2017). Many bacterial genomes also show a pronounced bias for genes to be on the leading strand of replication (Rocha and Danchin 2003; Rocha 2008). This is likely because expression from the lagging strand results in head-on collisions between DNA and RNA polymerases that can stall replication or abort transcription (Srivatsan et al. 2010; Merrikh et al. 2012). Indeed, when native or reporter genes are oriented against the dominant direction of transcription, they frequently suffer low expression or interfere with the expression of their neighbours (Bryant et al. 2014; Yeung et al. 2017; Ferrándiz et al. 2014). Thus, it is easy to see how horizontally transferred (HT) genes may similarly wind up incongruously expressed or disrupt the expression of nearby genes.

There are several lines of evidence that adaptive genome architecture has constrained the chromosomal topology of HGT. First, HT genes are more common on secondary chromosomes/chromids and plasmids, where essential and highly expressed genes are concomitantly rare (Rocha 2008; Couturier and Rocha 2006; Cooper et al. 2010). Such spatial segregation evidently minimizes disruption of evolved architecture on the main chromosome. Second, HT genes are often clustered (Dilthey and Lercher 2015; Touchon et al. 2009; Oliveira et al. 2017). In part, this reflects co-transfer of functionally co-dependent genes that would provide no benefit if transferred independently (Dilthey and Lercher 2015; Lawrence and Roth 1996). In part, however, it also reflects the existence of permissive zones or hotspots along the chromosome, which experience recurrent integration and high turnover (Oliveira et al. 2017). Permissive zones can originate or be reinforced through mechanistic integration biases, where the presence of integrases and/or recombinogenic sites (such as repeats and *dif* sites in *Escherichia coli*) facilitates acquisition of genetic material (Touchon et al. 2014). Third, regarding replication-related constraints, in multiple species, including *E. coli* and *Bacillus subtilis*, HT genes preferentially accumulate near the terminus (Rocha 2004; Zarei et al. 2013; Moszer et al. 1999; Touchon et al. 2014; Lawrence and Ochman 1998), perhaps to avoid deleteriously high expression near the origin. Analogously, on the linear chromosomes of Streptomyces *spp*., transposable elements chiefly populate the telomeric regions (Choulet et al. 2006). A recent aggregate analysis of HGT hotspots across 80 diverse bacterial taxa partly confirmed these initial observations: the frequency of hotspots containing prophages was found to increase along the origin-terminus (*ori-ter*) axis (Oliveira et al. 2017). However, hotspots lacking mobile element genes showed no such trend. This is arguably unexpected, given that replication is often considered a dominant force in defining bacterial genome organization. We therefore wondered whether topological biases may in fact exist but be obscured, for example, by aggregate analysis across bacterial clades with divergent organizational constraints or by a history of genomic rearrangements.

Here, we consider patterns of gene flow into a clade that includes Geobacillus, Parageobacillus, and Anoxygeobacillus *spp*., hereafter referred to as the GPA clade (Fig. 1A, Supplementary Table 1). Focusing on a single clade allows us to reconstruct HGT patterns with high confidence and high spatial and temporal resolution, which is difficult to replicate on a pan-bacterial scale. In addition, large-scale recombination events in Bacilli are dominated by symmetric inversions around the *ori-ter* axis (Repar and Warnecke 2017). This is useful for two reason: first, the preponderance of symmetric inversions indicates that genome architecture along the *ori-ter* axis is under strong constraint, suggesting we might also recover topological constraints on HGT. Second, the relationship between HGT and genome architecture is less likely to be obscured by asymmetric historical rearrangements. The results of our analysis, reported below, suggest that the topology of HGT in Geobacillus and other Bacilli reflects selection to minimize interference not only with fast growth but also with sporulation, as asymmetric chromosome confinement during sporulation imposes topological constraints on where HGT-driven innovation in early spore formation can occur.

**Figure 1.**
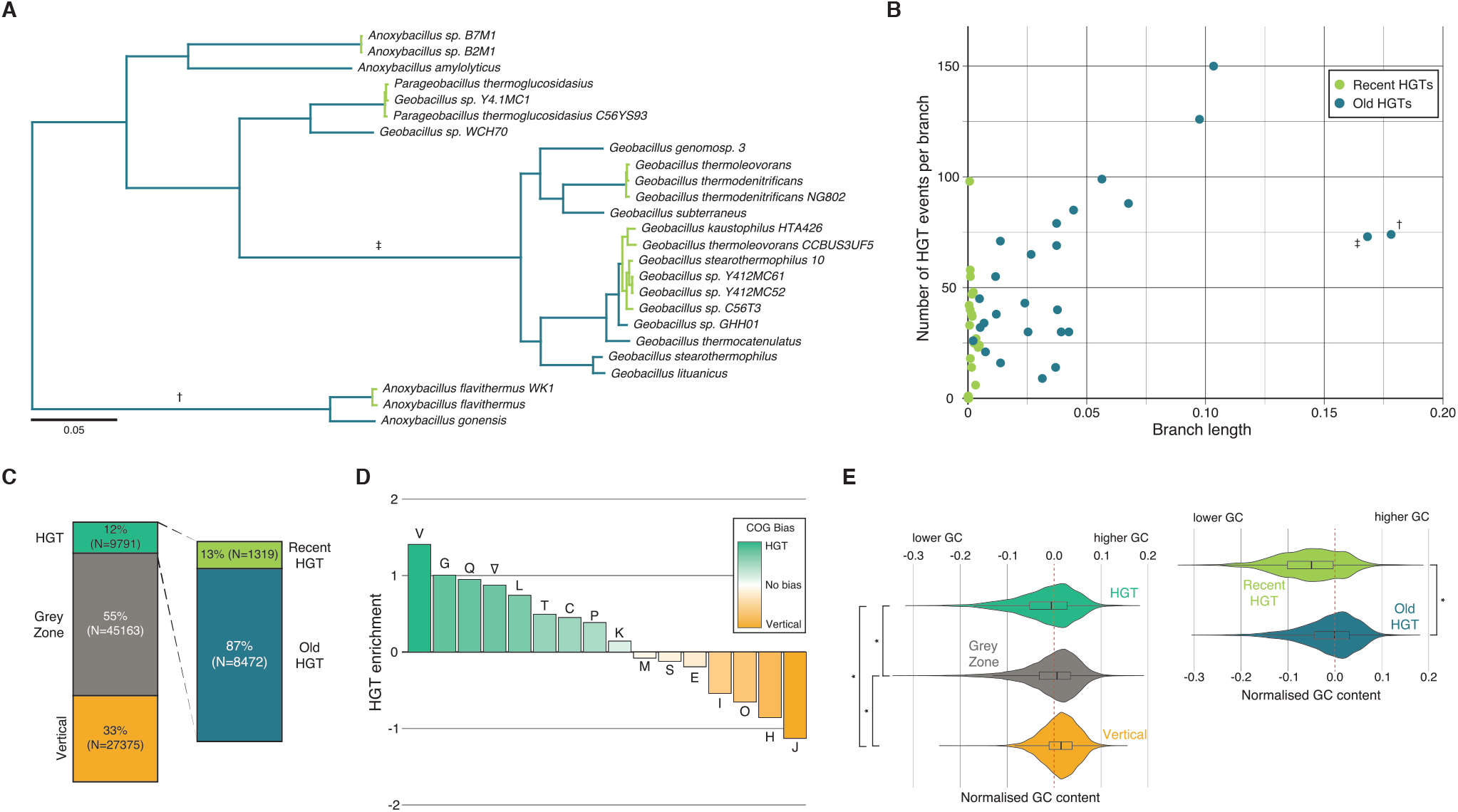
Global patterns of horizontal gene transfer into the GPA clade. (*A*) Reference GPA tree. The dashed grey line indicates the root, where the GPA tree is embedded in the larger reference tree. HGT events that occurred on light green branches were classified as recent. Dagger/double dagger symbols mark individual branches highlighted in panel (*B*). This panel illustrates the relationship between branch length and the number of inferred HGT events per branch across the GPA clade. *(C)* Absolute number (and proportion) of genes classified as HGT, vertical or belonging to the grey zone, as discussed in the main text. *(D)* Enrichment and depletion of COG functional classes amongst HT genes. The ∇ symbol indicates genes without an assigned COG functional class. Only COG classes with at least 50 associated genes are shown. *(E)* Distributions GC contents of HT/vertical/grey zone genes. GC contents are normalised to the mean GC content of all genes on a given chromosome to allow comparison across genomes with different GC contents. * P<0.001

## Results & Discussion

### Reconstructing horizontal gene transfers into the GPA clade

Prior analyses of the Geobacillus core and accessory genome indicated that HGT is common in this clade of bacteria (Studholme 2015; Bezuidt et al. 2016). However, explicit patterns of gene flow into this clade have not been characterized on a genome-wide scale. To do so, we first used a combined BLAST and Markov Clustering (MCL) approach to identify orthologous relationships between annotated proteins from 5073 fully assembled genomes, including 25 GPA genomes (see Methods, Supplementary Table 1). We then applied a phylogenetic approach to identify candidate HGT events. Principally, this involves evaluating the congruence between individual gene trees and a reference tree thought to reflect vertical inheritance (often referred to as the species tree). We reconstructed phylogenies for each orthologous group and used model-based reconciliation, implemented in Mowgli (Nguyen et al. 2013; Doyon et al. 2010), to a) identify incongruence and b) differentiate HGT from recurrent gene loss events, which can produce HGT-like phylogenetic signals. Full details of gene/species tree reconstruction and HGT inference procedures are provided in the Methods section.

Here, we should briefly highlight the following: First, the topology of the GPA part of the reference tree (Fig. 1A) is broadly consistent with branching patterns observed in prior analyses of conserved genes (Goh et al. 2014; Aliyu et al. 2016; Weng et al. 2009; Tourova et al. 2008; Coorevits et al. 2012; Studholme 2015; Zeigler 2005). Second, our principal aim was to identify high-confidence sets of HT and ‘vertical’ genes (defined as genes with an exclusively vertical evolutionary history in the GPA clade). To this end, rather than categorizing every gene as either horizontal or vertical, we conservatively defined a large ‘grey zone’ to include genes that are inconsistently predicted depending on parameter choices such as the penalty levied for calling an HGT event (versus loss) in Mowgli (see Methods). We also allocate putative transfers from within the Bacillaceae, the family in which GPA sits, to the grey zone (see Methods) to minimize spurious HGT calls and focus on transfers that are not homologous replacements. Note, however, that including HGT candidates from the grey zone in the analysis does not qualitatively affect results. The final set analysed below includes 27375 vertical genes and 2219 independent transfer events. Across 25 genomes, 9791 genes (11.9% of all GPA genes; Fig. 1C) are predicted as horizontally transferred into GPA, affecting 28.5% of orthologous groups (Supplementary Table 2). Given the stringent cut-offs applied here, and given that we exclude transfers from close relatives, this likely represents a lower-bound estimate.

The reconciliation-based approach predicts the phylogenetic branch along which a given gene was acquired. We find that the number of HGT events predicted per branch correlates well with the length of the branch (*r* = 0.58, *p* = 1.49e-05, Fig. 1B), supporting an approximately continuous model of HGT in this clade with no evidence of punctuated bursts of gene acquisition. There are two outlier branches: one (branch † in Fig. 1A) subtends a group of GPA genomes with elevated GC content (mean 52% versus 43%), compromising branch length as a proxy for time; the other (branch ‡ in Fig. 1A) subtends three taxa with reduced genome sizes compared to other GPA clade members (mean 2.82 Mb versus 3.60 Mb), suggesting a period of net loss of genetic material. For downstream analysis, we further classified 1319 genes, which occurred on short terminal branches, as recent transfers (versus 8472 older HT genes, see Methods and Fig. 1A/C), based on the reasoning that more recent events, not having experienced sustained purifying selection, might show different evolutionary dynamics (Hao and Golding 2006). Similar to studies in other prokaryotes, and in line with analyses of the Geobacillus core/accessory genome (Bezuidt et al. 2016), we find HGT events to be enriched for metabolic and defense-related genes and depleted for informational genes (Fig. 1D). Further, in agreement with others (Daubin and Perrière 2003), we find that HT genes are generally more AT-rich than vertical genes (Fig. 1E), irrespective of the GC content of the host genome (Supplementary Fig. 1)

### GPA chromosomes are characterized by large HGT-permissive and -restrictive zones

To evaluate HGT in the context of genome organization we considered the distribution of HT and vertical genes along GPA chromosomes. Normalising for genome length and looking across all GPA branches, we find three broad domains of HGT enrichment (Fig. 2A). HT genes were depleted in the immediate vicinity of the origin (“origin domain”, see Fig. 2C), but enriched either side of it (“near-origin domain”). A third domain of HGT enrichment spanned the terminus (“terminus domain”) whereas HT genes were depleted in broad stretches along the flanks (“flank domain”) of the replichores. The HGT pattern in Figure 2A is strikingly symmetrical. In part, this reflects the predominance of symmetric inversions in this clade (Repar and Warnecke 2017), i.e. genes on the left replichore in some genomes, will be on the right replichore following inversion, and vice versa, generating a symmetric pattern in aggregate analysis. However, the pattern is also evident in single-genome HGT enrichment profiles (Supplementary Fig. 2). While inspecting these individual profiles, we noticed that *Geobacillus sp. Y412MC61* and *A. amylolyticus* depart from the symmetric pattern (Supplementary Fig. 2). On closer inspection, it became evident that these genomes have undergone recent asymmetric inversions that shifted the terminus region, generating imbalanced replichore lengths. To facilitate pan-genome analysis of HGT in the context of constrained genome architecture, we excluded these two genomes from further analysis.

**Figure 2.**
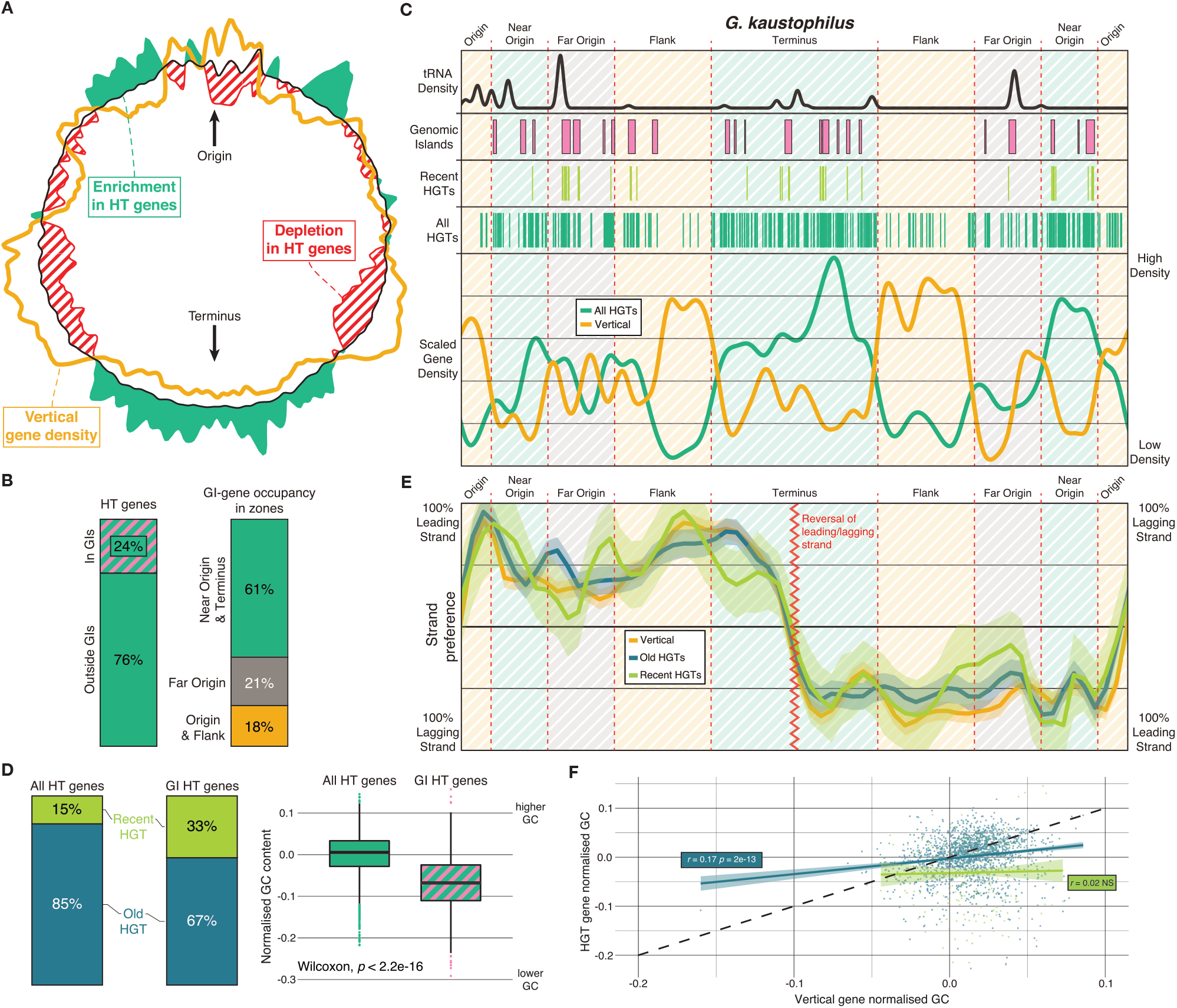
Topology of HGT into the GPA clade. (*A*) Density profiles illustrating the relative enrichment/depletion of HT(green/red) and vertical (yellow) genes across length-normalized GPA genomes. Overall gene density is given by the black line. (*B*) Representation across GPA genomes of HT genes inside and outside genomic islands (GI, left) and in different genomic zones (right) as defined in panel C. *(C)* Distribution of HT genes and other features along the *G. kaustophilus* genome. Zones were defined manually based on cross-genome enrichment profiles as shown in panel A (also see Fig. 3A). (D) GIs are enriched for recent HGT events compared to the genome-wide average and GI HT genes have a lower average GC content than HT genes outside GIs. *(E)* Relative leading/lagging strand enrichments for HT and vertical genes. *(F)* The GC content of old but not recent HT genes correlates with the GC content of their vertical neighbours, defined as vertical genes being in the same non-overlapping bin (each bin = 0.05% of the chromosome) in a given genome.

### Zones of HGT enrichment encompass and connect genomic islands

Having observed broad contiguous zones of HGT enrichment and depletion, we first asked how these zones relate to genomic islands (GIs). Genomic islands are generally considered to be relatively large (>10kb) HGT-derived DNA segments that differentiate closely related strains of bacteria. They harbour genes associated with mobility across genomes such as integrases and transposases, are characterized by divergent nucleotide content, are frequently inserted at tRNA genes or direct repeats, and typically carry genes thought to confer niche-specific advantages (e.g. antibiotic resistance) that lead to phenotypic/fitness differences between otherwise isogenic strains (Juhas et al. 2009). Considering genomic islands previously identified in 17 of our 23 genomes (Bertelli et al. 2017, see Methods), we find that the three zones encompass but are by no means synonymous with GIs. Of 6824 HT genes across the 17 genomes, only 1619 (24%) fall within predicted GIs (Fig. 2B). In other words, GIs sit within but do not define or delineate larger zones permissive for HGT. This is illustrated for *G. kaustophilus* in Figure 2C. GIs are enriched for more recent transfer events, marked by higher AT content (Fig. 2D), which is not unexpected given that nucleotide composition is used as a GI predictor (Bertelli et al. 2017). More notably, both recent and older transfers are strongly depleted in the flank and origin domains, suggesting rapid elimination from non-permissive zones and long-term stability of these zones (Fig. 2C). The distribution of genes on the leading and lagging strand further supports this model: in line with expectations (Touchon and Rocha 2016; Hao and Golding 2009), the majority of vertical GPA genes are located on the leading strand. But old and recent HT genes track the vertical pattern along the chromosome remarkably well. Assuming that the probability of initial integration to be approximately equal between the two strands, one would, in the absence of selection, have expected a more even leading/lagging strand distribution of recently transferred genes. We therefore suggest that HT genes, even if lowly expressed or non-essential, regularly disrupt adaptive local expression architecture (Fig. 2E), and are therefore quickly eliminated by selection, leading to conformity in strand preference between vertical and HT genes. In comparison, individual shortcomings that do not interfere with the expression of neighbouring genes might be tolerated for longer. As an example, we find no evidence for strong initial selection to conform to local GC content. Over the longer term, HT genes ameliorate to quantitatively mimic the GC content of their vertical neighbours (higher-GC vertical genes ending up with higher-GC HT companions), but recent transfers are equally AT rich, whether they integrate in a high or low GC neighbourhood (Fig. 2F).

### A link between horizontal gene transfer and chromosomal compartmentalization during sporulation

The observations above point to a compartmentalization of GPA genomes into broad HGT - permissive and -restrictive zones maintained by selection. Focusing on zones of HGT enrichment, we next asked why these zones are where they are. What explains the conspicuous enrichment of HGT events in the terminus and near-origin zones?

Prior work on gamma-proteobacteria in particular has highlighted the terminus as an exceptionally dynamic part of the genome. In *E. coli*, the terminus is enriched for HT genes (Rocha 2004; Lawrence and Ochman 1998), which are conditionally silenced by the nucleoid-associated protein H-NS (Zarei et al. 2013) and subject to high deletion rates (Koskiniemi et al. 2012; Louarn et al. 1991). This does not mean, of course, that there are no constraints on gene flow into the terminus region. For example, gene flow can be reduced if the incoming piece of DNA does not carry, in the right orientation, specific terminus-enriched motifs that are required for proper segregation (Hendrickson et al. 2018). Nonetheless, we were more surprised to see strong HGT enrichment relatively close to the origin of replication. It is not unusual to find pathogenicity islands located in the origin-proximal half of the replichore (Touchon et al. 2014; Westers et al. 2003), perhaps because their adaptive value hinges on high expression. But we demonstrated above that permissive zones transcend genomic islands and includes many older HT genes that are unlikely to fit a model where disruption of genome architecture is outweighed by transient benefits such as high expression of an antibiotic resistance gene. We therefore sought to better characterize the HT genes residing in different zones.

Considering broad functional categories as defined by Clusters of Orthologous Group (COG) functional classes, stark differences between zones emerge. First, although the origin zone is generally depleted for HT genes, it is the main new home for genes predicted to be involved in translation (COG J, Fig. 3A), tracking vertical enrichment patterns previously observed for fast-growing bacteria (Couturier and Rocha 2006). Second, the terminus zone is specifically enriched for genes associated with metabolic functions, including carbohydrate metabolism (COG G), energy production and conversion (COG C), amino acid metabolism (COG E), inorganic ion transport and metabolism (COG P) and secondary metabolite biosynthesis, transport and catabolism (COG Q, Fig. 3A). This is consistent with classic chromosomal compartmentalization into a fast-growth program encoded around the origin of replication and a slower, more metabolically challenging growth program that is executed in suboptimal conditions and draws more heavily on genes nearer the terminus. This partitioning is further supported by analyzing patterns of gene expression in different media. In the absence of large-scale expression data for GPA species, we considered publicly available expression data from *B. subtilis* strain 168 (Borkowski et al. 2016). We first identified orthologues of GPA-vertical genes on the *B. subtilis* chromosome, revealing extensive synteny (Fig. 3B). We then considered transcript levels in chromosomal context to find that the distribution of vertical genes along the chromosome is mirrored by average expression: areas of vertical enrichment near the origin and in the replichore flanks are on average more highly expressed (Fig. 3B). This is most pronounced during fast growth in casein hydrosylate medium supplemented with glucose (CHG). In poorer medium (M9SE, S), expression from genes around the terminus increases markedly, in line with condition-specific activation of metabolic genes. Rendering the topology of gene expression in this way highlights that chromosomal organization in Bacilli cannot be reduced to fast growth as the sole patterning force. It is true that genes close to the origin are more highly expressed, but rather than a gradual decrease in average expression towards the terminus, there is a sharp drop-off into the high-HGT near-origin zone, followed by modular-looking expression domains.

**Figure 3.**
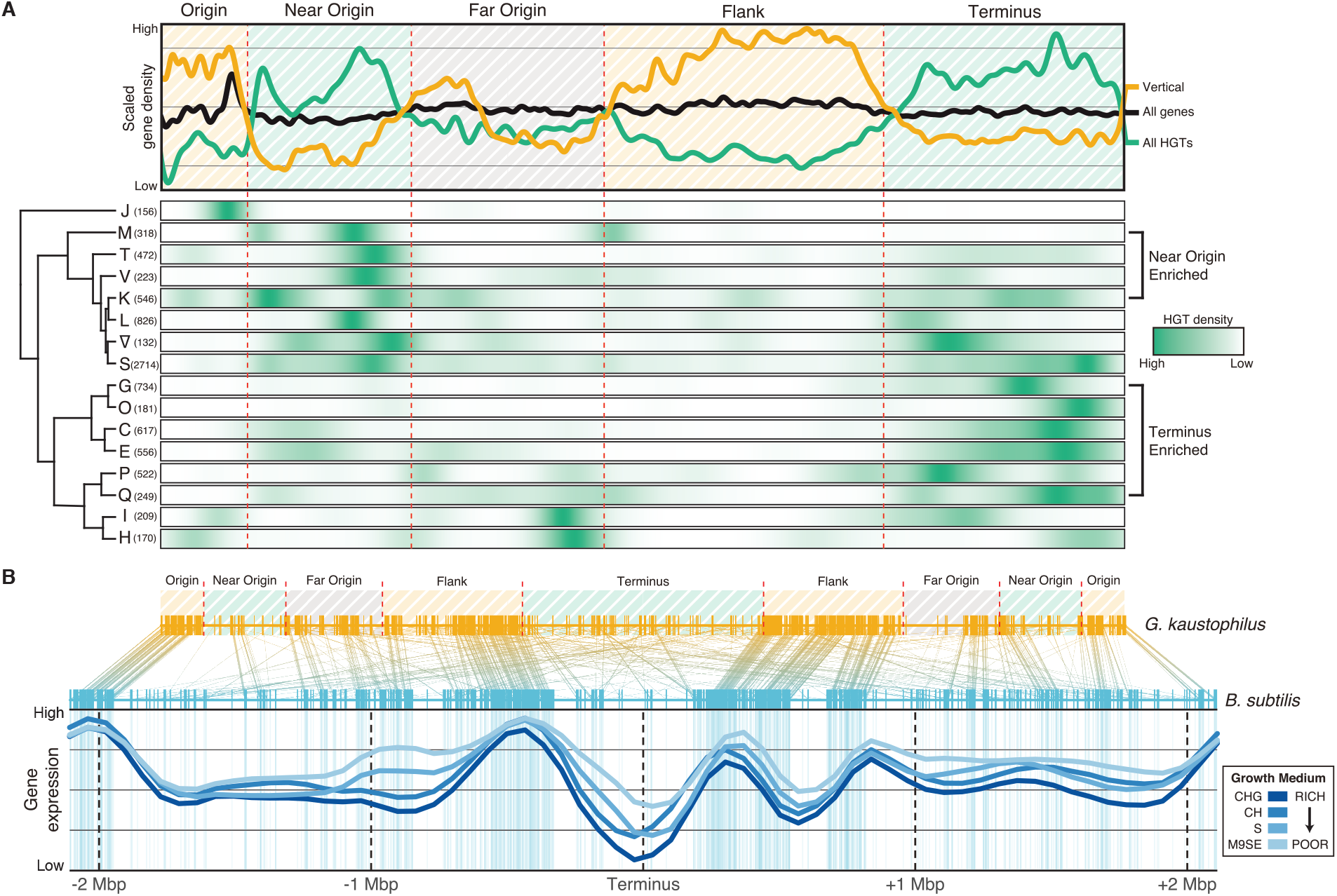
Spatio-functional patterns of gene flow into the GPA clade. (*A*) Global topological patterns of HGT across GPA chromosomes (top) are broken down by COG functional class (bottom). Left and right replichores have been combined for simplicity. HGT density is scaled by individual COG class rather than globally. Note in this context that high HGT densities for COG I and H in the Far Origin zone, and COG J in the origin zone are associated with relatively few genes, thus making little impression on overall HGT density. COG classes were hierarchically clustered based on the proportion of HGTs in the respective COG found in each zone. (*B*) High synteny of GPA-vertical genes in *G. kaustophilus*, chosen as a representative member of the GPA clade, and their orthologues in *B. subtilis* strain 168. Areas of high vertical gene density in *B. subtilis*, particularly in areas corresponding to GPA-defined Origin and Flank zones, are associated with higher average expression. Upregulation of expression in poorer media (lighter blue shades) is evident around the terminus and elsewhere. Thick blue lines represent LOESS regression fits between replicate-averaged expression values and gene midpoints.

Considering the near-origin zone, we find an enrichment for genes involved in transcription regulation (COG K) and signal transduction (COG T) as well as defence-related functions (COG V). Arguably the most noticeable enrichment, however, comes from genes related to cell wall/membrane biogenesis (COG M). 49.7% of HT genes that belong to this category cram into a region that comprises only 17% of the genome. Genes of unknown function (COG S/∇), as might be expected, are prevalent in both high-HGT zones (Fig. 3A).

We propose that the near-origin high-HGT zone is where it is and contains the types of genes that it does not because of constraints imposed by fast growth but by a developmental program characterized by lack of growth - sporulation. To see why, consider what happens at the early stages of sporulation, as reviewed for *B. subtilis* by Hilbert and Piggot (2004): after the sporulation program is initialized, the chromosome is duplicated and one copy localizes to each pole of the pre-divisional cell, a region around the origin being recognized by the RacA protein and tethered to the cell membrane (Fig. 4A). Then, the cell divides asymmetrically. A septum forms that establishes a mother cell and a prespore/forespore compartment. Importantly, septation traps approximately the first third of one of the chromosomes in the forespore (Frandsen et al. 1999). When projected onto a GPA genome, this corresponds remarkably well to the outer edge of the HGT-enriched near-origin zone. The remainder of the chromosome, along with the entire second copy, is at this point confined to the mother cell. Active translocation through the septum eventually reels the rest of the chromosome into the forespore, but for a critical period of early spore formation, genome dosage and expression in the two compartments are asymmetric (Frandsen et al. 1999). To contribute to gene expression in the very early forespore, before feeding channels to the mother cell have been established, genes need to be located on the first third of the chromosome. Indeed, there is a conspicuous cluster of genes controlled by σ^F^, the compartment-specific sigma factor that orchestrates gene expression in the early forespore (Losick and Stragier 1992), in the first third of the *B. subtilis* chromosome (Wang et al. 2006). In contrast, no such origin-proximal clustering is evident for targets of σ^G^, which replaces σ^F^ after the chromosome has fully entered the forespore and dosage equity has been restored (Wang et al. 2006). Notably, the same functional classes for which we find HGT enrichments in the near-origin zone in GPA are prominent targets in the σ^F^ regulon, as highlighted by Wang and colleagues. In the forespore-trapped first third of the *B. subtilis* chromosome, this includes the transcription regulators *yabT* and *rsfA* and a slew of genes that are either directly involved in spore morphogenesis or localized to the membrane (*spoIIQ, tuaF, ywnJ, pbpG, cydD*).

**Figure 4.**
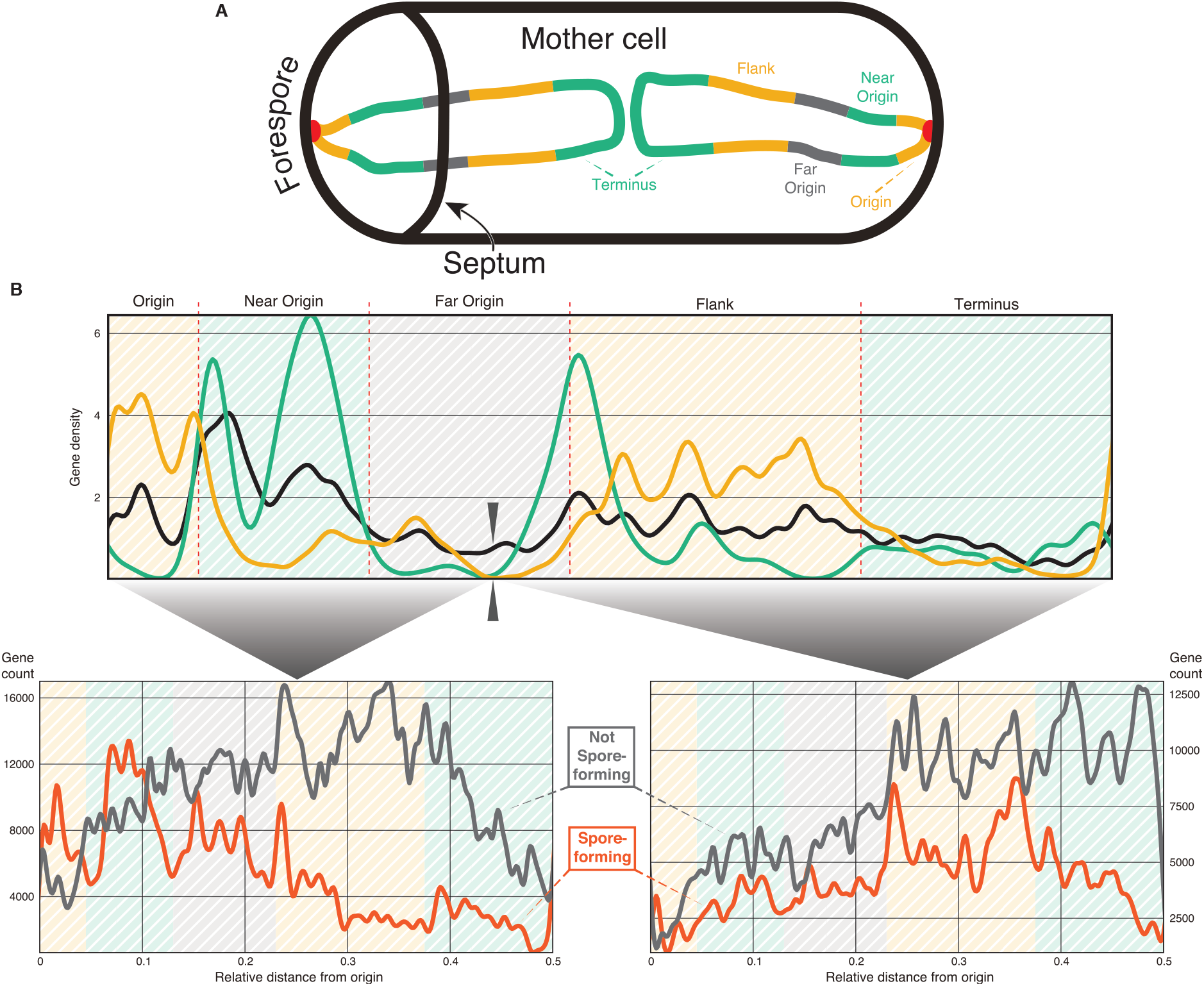
Topological patterns of HGT in light of spore formation (*A*) Schematic representation of HGT-enriched and –depleted zones in the forespore and mother cell during early spore formation. (*B*) The upper panel illustrates the relative density of COG M vertical (yellow) and HT (green) genes along GPA chromosomes. Left and right replichores have been combined for simplicity. The lower two panels show the topological distribution of orthologs of GPA COG M genes in spore-forming (red) and non-spore-forming (grey) Bacilli. Left: orthologues of those GPA genes that are located on the origin-proximal third of the chromosome. Right: orthologues of those GPA genes that are located in the terminus-proximal two-thirds.

To explore the link between spore formation, gene functional enrichment, and HGT further, we compared the location of genes of a given COG class in spore-forming and non-spore-forming Bacilli. We reasoned that, if relative location along the replichore is linked to the physical constraints of spore formation, we should see a stronger enrichment for COG M (membrane) genes in the near-origin zone in spore-forming than non-spore-forming Bacilli. This is indeed the case. Considering GPA COG M genes located on the first ~1/3 of the chromosome (first 38%, to coincide with the dip in overall gene density), we find that their orthologues are markedly enriched towards the origin, peaking in the near-origin zone, in spore-forming bacteria but not in organisms that lack the capacity to sporulate (Fig. 4B, lower left panel). Orthologues of those GPA COG M genes that are located in the latter two-thirds show no strong topological constraint in spore-forming bacteria (Fig. 4B, lower right panel). We find similar, albeit weaker trends, for COGs K and T (Supplementary Fig. 3). Based on these results, we suggest that asymmetric confinement of one chromosome during early forespore formation has affected the topological pattern of HGT into GPA *spp*. and likely other spore-forming Bacilli. To contribute to early development inside the forespore, genes need to be actively expressed from the first third of the chromosome. Real estate in the direct vicinity of the origin is limited and largely occupied by genes required for fast growth, so σ^F^-controlled early-acting genes cluster in the near-origin zone (Wang et al. 2006). Innovation in early spore morphogenesis via HGT faces the same topological constraints, concentrating HT genes in the same area. Membrane proteins might be particularly dependent on a prime slice of real estate in the forespore for two reasons: First, continued membrane synthesis is required for forespore growth after septation (López-Garrido et al. 2018), at a time when the opportunity for a maternal contribution is limited. Second, while some proteins can in principle be produced in the pre-divisional cell or provisioned from the mother cell via feeding channels, many membrane proteins rely on co-translational folding and/or insertion into the membrane and thus need to be produced in a time-sensitive manner in the relevant compartment. As a consequence, the distribution of membrane-associated genes, including those that arrived via HGT, mirrors the physical compartmentalization of the chromosome during sporulation. Note also that comparative genomic studies suggest that a contribution by HT genes to forespore development is likely: whereas the machinery responsible for the earliest stages of sporulation (initiation, septum formation, and asymmetric division) is highly conserved across Firmicutes, this is not the case for compartment-specific programs (Galperin et al. 2012; de Hoon et al. 2010); σ^F^-controlled genes in particular exhibit a narrow phylogenetic distribution suggestive of high turnover (de Hoon et al. 2010).

In summary, while HGT in bacteria is ubiquitous, gene flow into host chromosomes is not unrestricted. Our findings suggest that, in GPA and perhaps spore-forming bacteria more generally, fast growth and sporulation jointly constrain the chromosomal niches where HT genes can establish a foothold, both in terms of relative location along the replichore and in terms of leading/lagging strand orientation. As a result, gene flow has principally been directed into three broad contiguous zones: one around the terminus, preferentially housing condition-specific metabolic genes, and two either side of the origin of replication, where we encounter many genes with membrane function and/or localization. Comparison of older and more recent transfers suggest that zoning is rapidly enforced and, while individual genes come and go, zones themselves have existed in those locations for long periods of evolution. Permissive zones encapsulate many genomic islands, which we might think of not as isolated entities in a sea of vertical genes but rather as active volcanoes along a less visible undersea mountain range or as busy ports of entry into the permissive zone. In addition to helping us better understand the rules governing microbial evolution and innovation, our results (and approach) might also inform future genome engineering efforts. We focused on the GPA clade, in part, because their thermophilic lifestyle has made them attractive chassis for synthetic biology (Reeve et al. 2016; Cripps et al. 2009) and the topology of horizontal gene flow might provide clues as to where in the genome one should aim to introduce a given novel part. That chromosomal context affects (and is affected by) expression of an inserted construct is well recognized (Cardinale and Arkin 2012; Yeung et al. 2017). Yet, at a time when CRISPR/Cas technology has considerably simplified introducing blocks of DNA anywhere in the genome, predicting good chromosomal insertion sites remains a difficult challenge. Studying HGT patterns into a chassis of interest might help not only to identify permissive sites but also to provide more bespoke guidance for specific genes according to their expression requirements and cellular dependencies. Our study provides a template to guide future efforts in this direction.

## Methods

### Data acquisition

We downloaded 5073 bacterial genomes (assembly level: ‘complete genome’, Supplementary Table 1) from the NCBI Refseq database (https://www.ncbi.nlm.nih.gov/refseq/). Each of these genomes is associated with a unique taxonomic identifier (taxid). Where multiple assemblies were available for a single taxid, the most recent record was chosen. Translations of all annotated coding sequences were extracted from each genome using genbank_to_fasta.py (http://rocaplab.ocean.washington.edu/tools/genbank_to_fasta/). Entries lacking valid nonredundant protein accession identifiers (protIDs) were excluded. Where multiple protein sequences within a single genome shared a protID, the first instance in the file was retained, duplicates were transiently masked and re-added after orthologous gene family clustering (see below).

### Reconstructing GPA gene family histories

As our focus is exclusively on reconstructing gene flow *into* the GPA clade, gene family and phylogenetic reconstruction followed a GPA-centric approach. First, each of the 25 GPA proteomes was reciprocally BLASTed (blastp 2.2.28+) against every other proteome with the following parameters: -evalue 1E-10 -max_target_seqs 1 -max_hsps_per_subject 1 -seg yes - soft_masking true. To identify paralogs within individual genomes, GPA proteomes were BLASTed against themselves with parameters as above and all non-self BLAST hits were considered putative paralogs. Where the BLAST bitscore between two putative paralogs was greater than or equal to the maximum bitscore in the orthologous bitscores for either protein, the two proteins were considered true paralogs. All RBBH and paralog relationships were collated and subjected to clustering with MCL v. 14.137 (Enright et al. 2002) with the following parameters: --abc -I 2.0 -te 20 -scheme 7 --abc-neg-log10 -abc-tf ceil(200). Evalues served as graph edge weights. The clustering produced a set of orthologous gene families. At this stage, previously masked duplicate proteins were re-added to their gene families. Whenever predicted paralogs were split amongst two or more gene families (<1% of gene families), the gene families were removed from the analysis as this suggested mis-clustering. Finally, gene families with fewer than four proteins were removed as unsuitable for phylogenetic inference.

We then repeated the same BLAST-plus-MCL procedure for three additional BLAST Evalue thresholds (1E-150, 1E-100 and 1E-50, in addition to 1E-10). This was done for the following reason: Predicting HGTs using a phylogenetic approach requires sufficient orthologous context, but predicting HGTs *into a specific model clade* (rather than across the entire gene family) means that the context required to differentiate horizontal from vertical histories need not be comprehensive. For example, if GPA proteins in a sample gene family have 500 orthologues predicted at high homology (E-value = 1E-150) the gene tree likely provides sufficient phylogenetic context to predict whether the gene was transferred into GPA or vertically inherited. Including more orthologues predicted at a lower homology threshold (e.g. E-value = 1E-10) provides no obvious advantage since the local phylogenetic context of the GPA proteins will still be provided by those proteins with the highest homology, i.e. the 500 proteins at 1E-150. In such a scenario, a larger gene tree will not provide a better context for HGT prediction *into GPA* but will increase the computational cost of phylogenetic reconstruction and reconciliation. Defining gene families at different E-value thresholds, then, allows us to apply an ad hoc gene family reduction approach to optimise gene family size for downstream computation. We selected gene families at the most stringent threshold at which they contained at least 200 proteins, contained all GPA homologs identified at the most lenient 1E-10 base threshold, and did not contain GPA proteins other than those identified at the base threshold. Where any of the above criteria were violated, we defaulted to the base threshold gene family. This approach reduced the overall number of protein sequences in the analysis from 6547127 to 3789584, which was associated with significant computational cost savings.

The protein sequences belonging to each gene family were aligned using Clustal Omega v. 1.2 (Sievers et al. 2011) with default parameters. The alignments were then used as input for phylogenetic reconstruction using FastTree v. 2.1.8 (-gamma -boot 100) (Price et al. 2009). Although previous studies have highlighted that FastTree often produces worse gene trees than more computationally intensive alternatives such as RAxML (Stamatakis 2014) – see e.g. (Zhou et al. 2018) – we settled on FastTree as a faster, more scalable alternative after directly comparing performance (with regard to downstream classification into GPA-HT and–vertical genes) of FastTree and RAxML using a preliminary set of 19 GPA and 3953 other genomes. Predictions from RAxML and FastTree-derived phylogenies were highly congruent (Supplementary Fig. 4).

### Species tree reconstruction

As highlighted above, our focus was exclusively on predicting HGT events *into* the GPA clade, without concern for events in the wider phylogeny. For this purpose, our species tree needed to be representative (i.e. any taxon encountered in any of the gene trees needed to be present in the species tree) and particularly accurate for GPA species and their local phylogenetic context (operationally defined as Bacillaceae). Guided by these requirements, we constructed a representative species tree in two phases. First, we assembled a species tree with the coalescent-based method ASTRAL-II v. 5.5.9 (Mirarab and Warnow 2015) using the entire set of gene trees as input. For input into Mowgli (see below) the output tree, which represented all the 5070 taxa present in the underlying gene trees (two of the initial genomes were not present in any of the final reduced gene families), was mid-point rooted and rendered ultrametric. We then looked to confirm and – if necessary – refine the ASTRAL-II-derived topology for the Bacillaceae (and the GPA clade contained within) using an orthogonal supermatrix approach. To this end, we selected all proteomes from our processed dataset classified as Bacillaceae in the NCBI taxonomy database (N=215, including all 25 GPA proteomes), performed an all-versus-all BLAST, and clustered orthologous gene families at two E-value thresholds (1E-10, 1E-50) as described above. At both E-value thresholds, we selected those gene families in which the proteins were present in a single copy in every Bacillaceae genome (1-to-1 orthologues); these were then independently aligned using Muscle v. 3.8.31 (Edgar 2004) with default parameters and combined into a supermatrix. The phylogenies for both supermatrices (one for each initial E-value threshold) were reconstructed with both RAxML (parameters: -m PROTCATAUTO –p [random int] -x [random int] -N 100) and FastTree (parameters: -lg). In all cases, at each threshold, the topologies of the RAxML and FastTree reconstructions were identical. However, topologies differed marginally between thresholds: within the GPA clade, the two trees disagreed in the placement of *G. sp. GHH01*. To resolve this discrepancy, we turned to a prior reconstruction we performed on a broader set of GPA genomes (including those without completely assembled genomes, and so excluded from the rest of this work). This larger phylogeny strongly supported *G. sp. GHH01* as an outgroup to the *G. kaustophilus* and *G. sp. C56T3* subclades. This corresponded to the topology of the 1E-50 consensus tree, which we further use as the consensus Bacillaceae and GPA phylogeny.

We found the ASTRAL-II-derived and supermatrix-derived Bacillaceae phylogenies to be remarkably consistent. Within the GPA clade, we identified a single topological incongruence: the exact branching of *Geobacillus sp. GHH0*, which was corrected in the ASTRAL-II tree to conform to the supermatrix phylogeny. Discrepancies between ASTRAL-II and supermatrix topologies within the wider Bacillaceae were limited to closely related strains of the same species and did not warrant further corrections.

### Reconciliation

Gene evolutionary histories were reconstructed using *MowgliNNI* v. 2.0 (Nguyen et al. 2013) with individual gene trees and the corrected ASTRAL-II species tree as inputs. Reconciliation requires the provision of evolutionary cost parameters – the relative likelihoods of gene loss, duplication, and transfer in the evolutionary history of the gene, i.e. the major biological processes that might alternatively explain conflicting gene tree and species tree topologies. At a higher transfer:loss cost ratio, the reconciliation is more likely to predict a history dominated by multiple losses than by HGTs. Although it is often assumed that the probability of loss is higher than the probability of HGT, true costs will vary between gene families and between species within a gene family, so choosing appropriate relative costs is not straightforward. Even if parameters can be estimated from the data, overfitting becomes a concern where parameters are allowed to vary across the phylogeny. As a simpler alternative approach, we mitigated the effect of relying on arbitrary cost values by running the reconciliation for each gene tree with different transfer:loss cost ratios. Starting at a previously estimated cost relationship of: Loss = 1, Duplication = 2, Transfer = 3 (David and Alm 2011), we incrementally increased the transfer cost by 1, up to a transfer cost 6. We then selected only those gene trees in which a specific HGT event into GPA was robustly predicted at transfer costs of 3 and 4 and further checked whether downstream results remained robust for higher transfer costs, where the total number of events starts to decline. This strikes a balance between robust HGT prediction and losing too many – likely genuine – transfers because of prohibitively large transfer costs. Conversely, to be considered a vertical GPA gene, no inferred HGT was allowed to be present at any of the transfer costs used. Gene families with inconsistent predictions (between costs 3 and 4) regarding HGT into GPA were allocated to the grey zone. To further increase the stringency of HGT prediction, and avoid spurious HGT prediction due to gene tree reconstruction errors, we selected only those HGT events with predicted donors from outside the Bacillaceae. Finally, to be considered as a genuine HGT event, we required a given transfer to be predicted into the same branch of the GPA phylogeny across transfer:loss ratios.

### Gene function assignment

Gene functions (COG classes) were assigned using EggNOG 4.5.1 (Huerta-Cepas et al. 2016). All GPA proteins were individually queried using emapper.py against the EggNog bact_50 database. Of the 6726 orthologous gene families, 776 failed to find any orthologues within the EggNOG database and are designated as ∇ in Figures 1 and 3. Whenever a single protein belonged to multiple COG classes, all applicable COG classes were included.

### Assigning the origin of replication and left/right replichores

The precise location of replication origins is not known for many of the genomes used in this work, including many GPA genomes. This is true both with regard to experimental origin determination and computational predictions like those present in the frequently used DoriC database (Gao et al. 2013). For consistency, we therefore defined origin location as the first coding base of the *dnaA* gene and assigned left/right replichore accordingly (with the right replichore being downstream of *dnaA*). The *dnaA* gene had previously been found to be a good proxy for origin location (Mackiewicz et al. 2004) and is used as a key predictor in DoriC. We confirmed that *dnaA* location is indeed a good proxy by correlating origin locations as predicted across all genomes in DoriC with *dnaA* locations. Across bacteria, *dnaA* was within 1% of the genome length from the annotated origin in 88.6% of genomes and was immediately adjacent to the predicted origin in all GPA genomes present in DoriC.

### Genomic Island Predictions

Positions of GIs were downloaded from the IslandViewer 4 database (Bertelli et al. 2017), using the ‘all methods’ set. GI predictions were available for 19 of the 25 initial GPA genomes (17 of the 23 GPA genomes without a significant asymmetric rearrangement).

### Gene expression data

Microarray data from *B. subtilis* strain 168 grown in different media (Borkowski et al. 2016) were obtained from NCBI GEO (www.ncbi.nlm.nih.gov/geo/query/acc.cgi?acc=GSE78108). For visualization in Figure 3B, expression values were averaged across replicates in each condition.

### Spore-forming vs non-spore-forming Bacilli

All Bacilli genomes in our dataset were classified as either spore-forming or non-spore-forming based on prior family-level evidence for spore-formation capacity (Galperin 2013). All genomes within this work belonging to Alicyclobacillaceae, Bacillaceae, Paenibacillaceae, Sporolactobacillaceae and Thermoactinomycetaceae were considered as spore-forming. All genomes belonging to Listeriaceae, Staphylococcaceae, and the order Lactobacillales were considered non-spore-forming. We then excluded Bacillaceae, Sporolactobacillaceae and Thermoactinomycetaceae from further analysis to end up with a non-spore-forming set of genomes that is more closely related to the GPA clade than the spore-forming set. This was done to control for phylogenetic drag, i.e. the possibility of observing more GPA-like trends in a given set of genomes simply because those genomes happen to be more closely related. By taking the above steps, we conservatively stack the deck against this possibility. Finally, where multiple genomes corresponded to the same taxonomic species (see Supplementary Table 1), orthologue positions were averaged across all underlying genomes to avoid bias from highly sequenced taxa.

## Acknowledgments

The authors would like to thank members of the Warnecke and Ellis labs for feedback. AE is funded by an Imperial College Integrative Experimental and Computational Biology Studentship. TE is funded by EPSRC fellowship EP/M002306/1. TW receives core funding from the UK Medical Research Council.

## Author contributions

AE carried out all analyses. AE and TW designed the analyses and interpreted the data with input from TE. AE and TW conceived the project. All authors contributed to the writing of the manuscript.

